# On the scope and limitations of baker’s yeast as a model organism for studying human tissue-specific pathways

**DOI:** 10.1101/011858

**Authors:** Shahin Mohammadi, Baharak Saberidokht, Shankar Subramaniam, Ananth Grama

**Affiliations:** Department of Computer Sciences, Purdue University, West Lafayette IN 47904, USA; Department of Bioengineering, University of California at San Diego, La Jolla CA 92093, USA

## Abstract

Budding yeast, S. cerevisiae, has been used extensively as a model organism for studying cellular processes in evolutionarily distant species, including humans. However, different human tissues, while inheriting a similar genetic code, exhibit distinct anatomical and physiological properties. Specific biochemical processes and associated biomolecules that differentiate various tissues are not completely understood, neither is the extent to which a unicellular organism, such as yeast, can be used to model these processes within each tissue.

We propose a novel computational and statistical framework to systematically quantify the suitability of yeast as a model organism for different human tissues. We develop a computational method for dissecting the human interactome into tissue-specific cellular networks. Using these networks, we simultaneously partition the functional space of human genes, and their corresponding pathways, based on their conservation both across species and among different tissues. We study these subspaces in detail, and relate them to the overall similarity of each tissue with yeast.

Many complex disorders are driven by a coupling of housekeeping (universally expressed in all tissues) and tissue-selective (expressed only in specific tissues) dysregulated pathways. We show that human-specific subsets of tissue-selective genes are significantly associated with the onset and development of a number of pathologies. Consequently, they provide excellent candidates as drug targets for therapeutic interventions. We also present a novel tool that can be used to assess the suitability of the yeast model for studying tissue-specific physiology and pathophysiology in humans.

## Introduction

Budding yeast, *S. cerevisiae*, is widely used as an experimental system, due to its ease of manipulation in both haploid and diploid forms, and rapid growth compared to animal models. Coupled with the continuous development of new experimental methodologies for manipulating various aspects of its cellular machinery, it has served as the primary model organism for molecular and systems biology(1). Motivated by the availability of its full genome in 1996 as the first eukaryotic organism to be sequenced(2), an array of functional genomics tools emerged, including a comprehensive collection of yeast deletion mutants(3; 4), genome-wide over-expression libraries(5), and green fluorescent protein (GFP)-tagged yeast strains(6; 7). The maturity of yeast’s genetic and molecular toolbox has, in turn, positioned it as the primary platform for development of many high-throughput technologies, including transcriptome (8; 9; 10), proteome (11), and metabolome (12; 13) screens. These *-omic* datasets, all originally developed in yeast, aim to capture dynamic snapshots of the state of biomolecules during cellular activities. With the advent of “systems modeling”, a diverse set of methods have been devised to assay the interactions, both physical and functional, among different active entities in the cell, including protein-protein(14; 15; 16), protein-DNA(17; 18), and genetic(19; 20; 21) interactions. These interactions, also referred to as the *interactome*, embody a complex network of functional pathways that closely work together to modulate the cellular machinery. Comparative analysis of these pathways relies on network alignment methods, much the same way as sequence matching and alignments are used for individual genes and proteins. Network alignments use both the homology of genes, as well as their underlying interactions, to project functional pathways across different species(22; 23; 24; 25). These methods have been previously applied to detection of ortholog proteins, projection of functional pathways, and construction of phylogenetic trees.

Yeast and humans share a significant fraction of their functional pathways that control key aspects of eukaryotic cell biology, including the cell cycle (26), metabolism(27), programmed cell death(28; 29), protein folding, quality control and degradation(30), vesicular transport(31), and many key signaling pathways, such as mitogen-activated protein kinase (MAPK)(32; 33), target of rapamycin (TOR)(34), and insulin/IGF-I(35) signaling pathways. In the majority of cases, yeast has been the model organism in which these pathways were originally identified and studied. These conserved biochemical pathways drive cellular growth, division, trafficking, stress-response, and secretion, among others, all of which are known to be associated with various human pathologies. This explains the significant role for yeast as a model organism for human disorders(36; 37; 38). Yeast has contributed to our understanding of cancers(39; 40; 41) and neurodegenerative disorders(42; 43; 44). Having both chronological aging (amount of time cells survive in post-mitotic state) and replicative aging (number of times a cell can divide before senescence occurs), yeast is also used extensively as a model organism in aging research. It has contributed to the identification of, arguably, more human aging genes than any other model organism(45).

Depending on the conservation of the underlying pathways, there are two main approaches to studying them in yeast. It has been estimated that, out of 2,271 known disease-associated genes, 526 genes (~ 23%) have a close ortholog in the yeast genome, spanning 1 out of every 10 yeast genes(46). For these orthologous pairs of disease-associated genes, we can directly increase the gene dosage of the endogenous yeast protein by using overexpression plasmids, or decrease it, through either gene knockout or knockdown experiments, in order to study gain- or loss-of-function phenotypes, respectively. A key challenge in phenotypic screens is that disrupting genes, even when they have close molecular functions, can result in characteristically different organism-level phenotypes. *Phenologs*, defined as phenotypes that are related by the orthology of their associated genes, have been proposed to address this specific problem(47). A recent example of such an approach is the successful identification of a highly conserved regulatory complex implicated in human leukemia(48). This complex, named COMPASS (Complex of Proteins Associated with Set1), was originally identified by studying protein interactions of the yeast Set1 protein, which is the ortholog of the human mixed-lineage leukemia (MLL) gene, and years later was shown to be conserved from yeast to fruit flies to humans. On the other hand, if the disease-associated gene(s) in humans does not have close orthologs in yeast, heterologous expression of the human disease-gene in yeast, also referred to as *“humanized yeast”*, can be used to uncover conserved protein interactions and their context, to shed light on the molecular mechanisms of disease development and progression. For the majority of disease-genes with known yeast orthologs, heterologous expression of the mammalian gene is functional in yeast and can compensate for the loss-of-function phenotype in yeast deletion strains(1). This approach has already been used to construct humanized yeast model cells to study cancers(39), apoptosis-related diseases(49), mitochondrial disorders(50), and neurodegenerative diseases(43). Perhaps one of the more encouraging examples is the very recent discovery of a new compound, N-aryl benzimidazole (NAB), which strongly protects cells from *α*-synuclein toxicity in the humanized yeast model of Parkinson’s disease(51). In a follow-up study, they tested an analog of the NAB compound in the induced pluripotent stem (iPS) cells generated from the neuron samples of Parkinson’s patients with *α*-synuclein mutations. They observed that the same compound can reverse the toxic effects of *α*-synuclein aggregation in neuron cells(52). Using this combined phenotypic screening, instead of the traditional target-based approach, they were not only able to discover a key compound targeting similar conserved pathways in yeast and humans, but also uncover the molecular network that alleviates the toxic effects of *α*-synuclein. These humanized yeast models have also been used to study human genetic variations(53).

Various successful instances of target identification, drug discovery, and disease network reconstruction using humanized yeast models have established its role as a model system for studying human disorders. When coupled with more physiologically relevant model organisms to cross-validate predictions, yeast can provide a simple yet powerful first-line tool for large-scale genetic and chemical screening(43; 41). However, as a unicellular model organism, yeast fails to capture organism-level phenotypes that emerge from inter-cellular interactions. Perhaps, more importantly, it is unclear how effectively it can capture tissue-specific elements that make a tissue uniquely susceptible to disease. All human tissues inherit the same genetic code, but they exhibit unique functional and anatomical characteristics. Similar sets of molecular perturbations can cause different tissue-specific pathologies given the network context in which the perturbation takes place. For example, disruption of energy metabolism can contribute to the development of neurodegenerative disorders, such as Alzheimer’s, in the nervous system, while causing cardiomyopathies in muscle tissues(54). These context-dependent phenotypes are driven by genes that are specifically or preferentially expressed in one or a set of biologically relevant tissue types, also known as *tissue-specific* and *tissue-selective* genes, respectively. Disease genes, and their corresponding protein complexes, have significant tendencies to selectively express in tissues where defects cause pathology(55; 56). How tissue-selective pathways drive tissue-specific physiology and pathophysiology is not completely understood; neither is the extent to which we can use yeast as an effective model organism to study these pathways.

We propose a quantitative framework to systematically measure the suitability of yeast as a model organism for different human tissues. Our framework is grounded in a novel statistical model for effectively assessing the similarity of each tissue with yeast, considering both expressed genes and their underlying physical interactions as a part of functional pathways. To understand the organization of human tissues, we present a computational approach for partitioning the functional space of human proteins and their interactions based on their conservation both across species and among different tissues. Using this methodology, we identify a set of *core genes*, defined as the subset of the most conserved housekeeping genes between humans and yeast. These core genes are not only responsible for many of the fundamental cellular processes, including translation, protein targeting, ribosome biogenesis, and mRNA degradation, but also show significant enrichment in terms of viral infectious pathways. On the other hand, human-specific housekeeping genes are primarily involved in cell-to-cell communication and anatomical structure development, with the exception of mitochondrial complex I, which is also human-specific. Next, we identify comprehensive sets of tissue-selective functions that contribute the most to the computed overall similarity of each tissue with yeast. These conserved, tissue-selective pathways provide a comprehensive catalog for which yeast can be used as an effective model organism. Conversely, human-specific, tissue-selective genes show the highest correlation with tissue-specific pathologies and their functional enrichment resembles highly specific pathways that drive normal physiology of tissues.

Comparative analysis of yeast and human tissues to construct conserved and non-conserved functional tissue-specific networks can be used to elucidate molecular/ functional mechanisms underlying dysfunction. Moreover, it sheds light on the suitability of the yeast model for the specific tissue/ pathology. In cases where suitability of yeast can be established, through conservation of tissue-specific pathways in yeast, it can serve as an experimental model for further investigations of new biomarkers, as well as pharmacological and genetic interventions.

## Results and discussion

In this section, we present our comparative framework to investigate the scope and limitations of yeast as a model organism for studying tissue-specific biology in humans. Figure 1 illustrates the high-level summary of our study design. We start by aligning human tissue-specific networks with the yeast interactome. We couple the alignment module with a novel statistical model to assess the significance of network alignments and use it to infer the respective similarity/dissimilarity of human tissue-specific networks with their mapped counterparts in yeast. Using a network of tissue-tissue similarities, we show that our alignment *p*-values fall within coherent groups of tissues that exhibit consistent characteristics. We use this network to identify four such groups of tissues/ cell-types. Furthermore, we partition both housekeeping and tissue-selective subsets of human genes separately into the conserved and human-specific subsections. We provide extensive validation for the selective genes with respect to blood cells and brain tissues. Figure 2 illustrates the overall partitioning of these genes and their relative subsets. We provide an in-depth analysis of each of these subsets, and show that while conserved subsets can provide the *safe zone* that yeast can be used as an ideal model organism, the human-specific subset can shed light on the *shadowed subspace* of human genome in yeast and can provide future directions for constructing humanized yeast models.

**Fig. 1.**
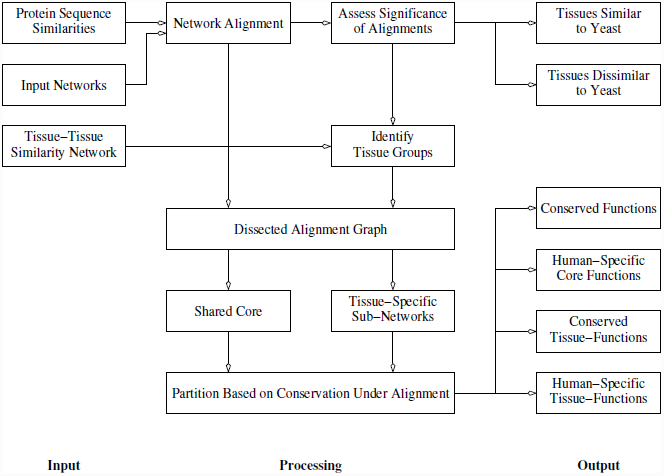
Workflow summary. Main components of the analysis framework proposed in this paper. Each intermediate processing step is further discussed in details in separate subsections.

**Fig. 2.**
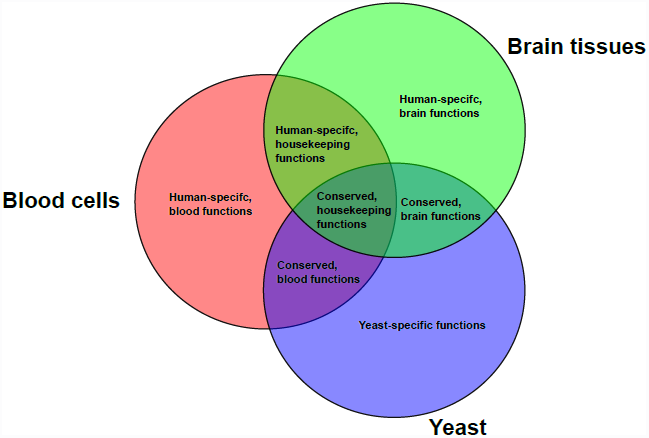
Functional classification of human genes. A high-level summary of gene classification performed in this study.

### Aligning yeast interactome with human tissue-specific networks

The *global* human interactome represents a static snapshot of potential physical interactions that *can* occur between pairs of proteins. However, it does not provide any information regarding the spatiotemporal characteristics of the actual protein interactions. These interactions have to be complemented with a dynamic *context*, such as expression measurements, to help interpret cellular rewiring under different conditions.

(57) overlaid the mRNA expression level of each transcript (transcriptome) in different human tissues(58) on top of the *global* human interactome, integrated from 21 PPI databases, and constructed a set of 79 reference tissue-specific networks. We adopt these networks and align each one of them separately to the yeast interactome that we constructed from the BioGRID database.

In order to compare these human tissue-specific networks with the yeast interactome, considering both the sequence similarity of proteins and the topology of their interactions, we employ a recently proposed sparse network alignment method, based on the Belief Propagation (BP) approach. This method is described in the Materials and methods section(59).

Genes, and their corresponding proteins, do not function in isolation; they form a complex network of interactions among coupled biochemical pathways in order to perform their role(s) in modulating cellular machinery. Moreover, each protein may be involved in multiple pathways to perform a diverse set of functions. Using a network alignment approach to project these pathways across species allows us to not only consider their first-order dynamics, through co-expression of homologous protein pairs, but also the context in which they are expressed.

To construct the state space of potential homologous pairs, we align all protein sequences in human and yeast and pre-filter hits with sequence similarity *E*-values greater than 10. For genes with multiple protein isoforms we only store the most significant hit. Using these sequence-level homologies, we construct a matrix *L* that encodes pairwise sequence similarities between yeast and human proteins. Entries in matrix *L* can be viewed as edge weights for a bipartite graph connecting human genes on one side, and the yeast genes, on the other side. We use this matrix to restrict the search space of the BP network alignment method (please see Supplementary Methods for details on *E*-value normalization and Materials and Methods section for BP alignment method).

Parameters *α* and *β*(= 1−*α*) control the relative weight of sequence similarity (scaled by *α*) as compared to topological conservation (scaled by *β*) in the BP network alignment. Using a set of preliminary simulations aligning the global human interactome with its tissue-specific sub-networks, for which we have the *true* alignment, with various choices of *α* in the range of 0.1 to 0.9, we identify the choices of 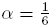 and 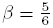 to perform the best in our experiments. We use the same set of parameters to align each tissue-specific network with the yeast interactome, as it provides a balanced contribution from sequence similarities and the number of conserved edges. The final set of all alignments is available for download as Additional file 1.

### Investigating roles of housekeeping genes and their conservation across species

Housekeeping genes comprise a subset of human genes that are universally expressed across all tissues and are responsible for maintaining core cellular functions needed by all tissues, including translation, RNA processing, intracellular transport, and energy metabolism(60; 61; 62). These genes are under stronger selective pressure, compared to tissue-specific genes, and evolve more slowly(63). As such, we expect to see a higher level of conservation among human housekeeping genes compared with yeast genes. We refer to the most conserved subset of housekeeping genes between humans and yeast, computed using network alignment of tissues-specific networks with the yeast network, as the *core genes*.

We identify a gene as housekeeping if it is expressed in *all* 79 tissues. We identify a total of 1,540 genes that constitute the shared section of human tissue-specific networks. These genes, while having similar set of interactions among each other, are connected differently to the set of tissue-selective genes.

Using the alignment partners of all housekeeping genes in the yeast interactome, we construct an alignment consistency table of size 1, 540 × 79, which summarizes the network alignments over the shared subsection of tissue-specific networks. Then, we use the majority voting method to classify housekeeping genes as *core*, which are conserved in yeast, *human-specific*, which are consistently unaligned across human tissues, and *unclassified*, for which we do not have enough evidence to classify it as either one of the former cases.

Network alignments are noisy and contain both false-positive (defined as aligned pairs that are not functionally related), as well as false-negatives (pairs of functional orthologs that are missed in the alignment). These errors can come from different sources, including gene expression data (node errors), interactome (edge errors), or the alignment procedure (mapping errors). We propose a method based on majority voting across different alignments to (partially) account for these errors. Given a set of network alignments, we consider a pair of entities consistently aligned (either matched or unmatched) if they are consistent in at least 100 * *τ*% of alignments in the set. The parameter *τ*, called the *consensus rate*, determines the level of accepted disagreement among different alignments. A higher value of consensus rate increases the precision of the method at the cost of decreased sensitivity. In order to select the optimal consensus rate parameter, we tried values in range [0.5 − 1.0] with increments of 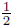. We identified the parameter choice of *τ* = 0.9, equivalent to 90% agreement among aligned tissues, to perform the best in classifying human-specific and conserved genes, while keeping the sets well-separated. Using this approach, we were able to tri-partition 1,540 housekeeping genes into 595 conserved, 441 human-specific, and 504 unclassified genes, respectively. The complete list of these genes is available for download as Additional file 1.

In order to investigate the conserved sub-network of core genes, we construct their alignment graph as the Kronecker product of the subgraph induced by core genes in the human interactome and its corresponding aligned subgraph in yeast. Conserved edges in this network correspond to interologs, i.e., orthologous pairs of interacting proteins between yeast and human(64). The final alignment graph of the core housekeeping genes is available for download as Additional file 1.

Figure 3 shows the largest connected component of this constructed alignment graph. We applied the MCODE(65) network clustering algorithm on this graph to identify highly interconnected regions corresponding to putative protein complexes. We identified five main clusters, which are color-coded on the alignment graph, and are shown separately on the adjacent panels. Ribosome is the largest, central cluster identified in the alignment graph of core genes, and together with proteasome and spliceosome, constitutes the three most conserved complexes in the alignment graph. This complex is heavily interconnected to the eIFs, to modulate eukaryotic translation initiation, as well as proteasome, which controls protein degradation. Collectively, these complexes regulate protein turnover and maintain a balance between synthesis, maturation, and degradation of cellular proteins.

**Fig. 3.**
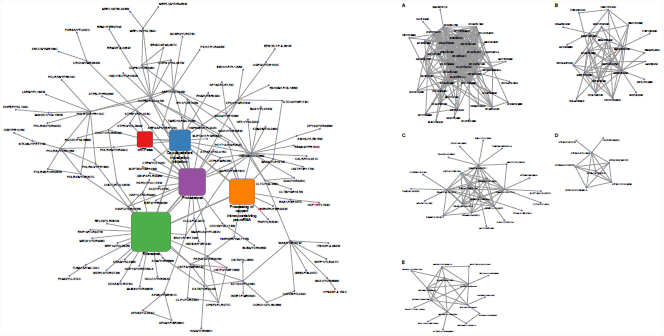
Alignment graph of core human genes. Conserved edges in the alignment graph of core housekeeping genes, which correspond to the “interologs,” i.e. orthologous pairs of interacting proteins between yeast and human. Five main protein clusters, identified as dense regions of interaction in the alignment graph, are marked accordingly and annotated with their dominant functional annotation as follows: **A** Ribosome, **B** Processing of capped intron-containing pre-mRNA, **C** Proteasome, **D** vATPase, **E** Cap-dependent translation initiation.

In order to further analyze the functional roles of these housekeeping genes, we use the g:Profiler(66) R package to identify highly over-represented terms. Among functional classes, we focus on the gene ontology (GO) biological processes, excluding electronic annotations, KEGG pathways, and CORUM protein complexes to provide a diverse set of functional roles. We use the Benjamini-Hochberg procedure to control for false-discovery rate (FDR), with *p*-value threshold of *α* = 0.05, and eliminate all enriched terms with more than 500 genes to prune overly generic terms. Using this procedure, we identify enriched functional terms for both core and human-specific subsets of housekeeping genes. The complete list of enriched functions for different classes of housekeeping genes is available for download as Additional file 1.

We manually group the most significant terms (*p*-value ≤ 10^−10^) in core genes, which results in five main functional classes, namely ribosome biogenesis, translation, protein targeting, RNA splicing, and mRNA surveillance. First, we observe a one-to-one mapping between enriched terms and identified putative complexes corresponding to translation initiation (*p*-value = 7.1*10^−17^) and ribosome (*p*-value = 5.97 * 10^−11^). In addition, translation termination and elongation are also enriched with decreasing levels of significance. Moreover, these processes are tightly linked to SRP-dependent co-translational protein targeting (*p*-value = 2.7 * 10^−15^). This, in turn, suggests protein synthesis as one of the most conserved aspects of eukaryotic cells. Next, we note that both mRNA splicing (*p*-value = 7.04*10^−10^) and nonsense-mediated decay (*p*-value = 4.66*10^−16^) are also enriched among the most significant functional terms, which supports our earlier hypothesis related to the role of splicesome in the alignment graph of core genes. Finally, we find that the most significant functional term, as well as a few other related terms, are involved in viral infection, which suggests that (a subset of the) core genes provides a *viral gateway* to mammalian cells. This can be explained in light of two facts: i) viral organisms rely on the host machinery for their key cellular functions, and ii) housekeeping genes are more ancient, compared to tissue-selective genes, and core genes provide the most conserved subset of these housekeeping genes. As such, these genes may contain more conserved protein interaction domains and be structurally more “familiar” as interacting partners for the viral proteins and provide ideal candidates for predicting host-pathogen protein interactions.

Next, we perform a similar procedure for the human-specific housekeeping genes. This subset, unlike core genes, is mostly enriched with terms related to anatomical structure development and proximal cell-to-cell communication (paracrine signaling), with the exception of complex I of the electron transport chain, which is the strongest identified term. This NADH-quinone oxidoreductase is the largest of the five enzyme complexes in the respiratory chain of mammalian cells. However, this complex is not present in yeast cells and has been replaced with a single subunit NADH dehydrogenase encoded by gene NDI1. Impairment of complex I has been associated with various human disorders, including Parkinson’s and Huntington’s disease. Transfecting complex I-defective cells with yeast NDI1 as a therapeutic agent has been proposed as a successful approach to rescue complex I defects(67; 68). This technique, also known as *NDI1 therapy*, opens up whole new ways in which yeast can contribute to the research and development on human diseases: not only yeast can be used as a model organism, but also can provide candidates that can be used for gene therapy in mammalian cells.

A key observation here is that the human-specific subset of housekeeping genes is not only associated with fewer functional terms, but is also less significantly associated with these terms. This effect can be attributed to two factors. First, we note that some of the genes predicted to be human-specific might be an artifact of the method. For example, the belief propagation (BP) method enforces sequence similarity as a necessary, but not sufficient, condition for a pair of genes to be aligned, which means that any human gene with no sequence similarity to yeast genes will not be aligned, resulting in genes being artificially classified as human-specific. Second, and more importantly, a majority of functional annotations for human genes are initially attributed in other species, specially yeast, and transferred across ortholog groups. Based on our construction, human-specific genes are defined as the subset of housekeeping genes with no orthology with yeast. As such, it can be expected that these genes span the *shadowed subspace* of the functional space of human genes that is under-annotated.

### Quantifying similarity of human tissues with yeast

Housekeeping genes are shared across all human tissues and cell types. They provide a conserved set of functions that are fundamental to cellular homeostasis. However, these genes do not provide direct insight into how different tissues utilize these key functions to exhibit their dynamic, tissue-specific characteristics. To assess the similarity of each tissue with yeast, we propose a novel statistical model, called *tissue-specific random model* (*TRAM*), which takes into account the ubiquitous nature of housekeeping genes and mimics the topological structure of tissue-specific networks (please see Materials and Methods section for the details of the random model).

We use the alignment score of each tissue-yeast pair as the objective function. To asses the significance of each alignment score, we use a Monte Carlo simulation method to sample from the underlying probability distribution of alignment scores.

For each tissue-specific network, we sample *k*_𝓡_ = 10, 000 pseudo-random tissues of the same size from TRAM, separately align them with the yeast interactome, and compute the number of conserved edges and sequence similarity of aligned protein pairs as alignment statistics, in order to compute the empirical *p*-values. For each network alignment, we compute a *topological*, a *homological* (sequence-based), and a *mixed* (alignment score) *p*-value. Additionally, we use cases in which alignment quality is significantly better in the original tissue alignment, both in terms of sequence and topology, to quantify an *upper bound* on the alignment *p*-values. Conversely, cases in which both of these measures are improved in the random samples can be used to define a *lower bound* on the alignment *p*-value. The final table of alignment *p*-values is available for download as Additional file 1.

First, we note that all tissues with significant *mixed p*-values also have both significant topological and homological (sequence-based) *p*-values. For a majority of tissues with insignificant *mixed p*-values, we still observe significant homological, but insignificant topological *p*-values. We summarize the most and the least similar tissues to yeast by applying a threshold of *α_l_* = *α_u_* = 10^−2^ to the *p*-value upper and lower bounds, respectively. There are a total of 23/79 tissues that have Δ*_𝓡_* ≤ 10^−2^ (*p*-value upper bound), listed in Table 1, which show the most significant similarity to the yeast interactome. Among these, blood cells show coherent high significance, with not even a single instance from 10, 000 samples having either the alignment weight or the overlap of the random sample exceeding the original alignment. Similarly, blood/immune cell lines consistently show significant alignment *p*-values. Male reproductive tissues also show a strong similarity to yeast cells. Conversely, there are 19/79 tissues with 10^−2^ < *δ_𝓡_* (*p*-value lower bound), which show the least significant similarity to yeast. Among these tissues, listed in Table 2, ganglion tissues consistently show the least similarity to yeast. An interesting observation is that tissues and cell types at either end of the table (either the most or the least similar) usually have very high confidence values, i.e., both their topology and homology *p*-values are consistent.

**Table 1.** Tissues with the most significant similarity to the yeast interactome.

| Name | pval lower bound | overall pval | pval upper bound | confidence |
| --- | --- | --- | --- | --- |
| Myeloid Cells | < 1.00e-04 | < 1.00e-04 | < 1.00e-04 | 1 |
| Monocytes | < 1.00e-04 | < 1.00e-04 | < 1.00e-04 | 1 |
| Dendritic Cells | < 1.00e-04 | < 1.00e-04 | < 1.00e-04 | 1 |
| NK Cells | < 1.00e-04 | < 1.00e-04 | < 1.00e-04 | 1 |
| T-Helper Cells | < 1.00e-04 | < 1.00e-04 | < 1.00e-04 | 1 |
| Cytotoxic T-Cells | < 1.00e-04 | < 1.00e-04 | < 1.00e-04 | 1 |
| B-Cells | < 1.00e-04 | < 1.00e-04 | < 1.00e-04 | 1 |
| Endothelial | < 1.00e-04 | < 1.00e-04 | < 1.00e-04 | 1 |
| Hematopoietic Stem Cells | < 1.00e-04 | < 1.00e-04 | < 1.00e-04 | 1 |
| MOLT-4 | < 1.00e-04 | < 1.00e-04 | < 1.00e-04 | 1 |
| B Lymphoblasts | < 1.00e-04 | < 1.00e-04 | < 1.00e-04 | 1 |
| HL-60 | < 1.00e-04 | < 1.00e-04 | < 1.00e-04 | 1 |
| K-562 | < 1.00e-04 | < 1.00e-04 | < 1.00e-04 | 1 |
| Early Erythroid | < 1.00e-04 | < 1.00e-04 | < 1.00e-04 | 1 |
| Bronchial Epithelial Cells | < 1.00e-04 | < 1.00e-04 | 0.0002 | 0.9998 |
| Colorectal Adenocarcinoma | < 1.00e-04 | < 1.00e-04 | 0.0004 | 0.9996 |
| Daudi | < 1.00e-04 | < 1.00e-04 | 0.0009 | 0.9991 |
| Testis Seminiferous Tubule | < 1.00e-04 | < 1.00e-04 | 0.0012 | 0.9988 |
| Smooth Muscle | < 1.00e-04 | < 1.00e-04 | 0.0016 | 0.9984 |
| Blood (Whole) | < 1.00e-04 | < 1.00e-04 | 0.0053 | 0.9947 |
| Thymus | < 1.00e-04 | 0.0001 | 0.0062 | 0.9938 |
| Testis Interstitial | < 1.00e-04 | 0.0004 | 0.0086 | 0.9914 |

**Table 2.**
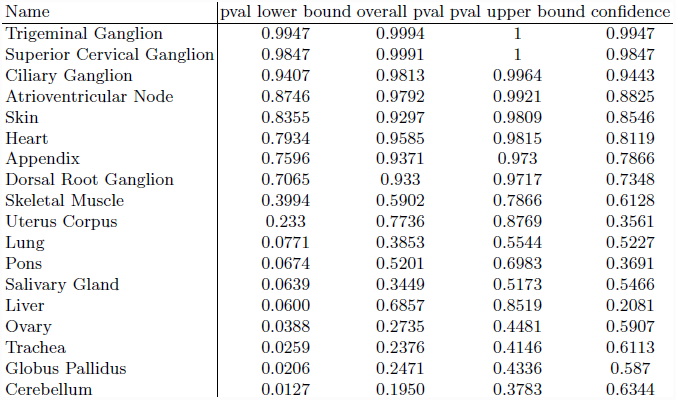
Tissues with the least significant similarity to the yeast interactome.

### Identifying groups of coherent tissues

Next, we investigate the correlation between the similarity of human tissues among each other and how it relates to their corresponding alignment *p*-values with yeast in order to better understand the transitivity of this relationship. We expect that similar tissues should exhibit consistent alignment *p*-values, resulting in groups of homogenous tissues with coherent alignments scores.

To this end, we first construct a network of tissue-tissue similarities (TTSN) using the global transcriptome of human tissues from the GNF Gene Atlas, including 44,775 human transcripts covering both known, as well as predicted and poorly characterized genes. For each pair of tissues/ cell types, we compute a similarity score using the Pearson correlation of their transcriptional signatures and use the 90 percentile of similarity scores to select the most similar pairs. We annotate each node in the TTSN with its corresponding alignment *p*-value as a measure of similarity with the yeast interactome. This meta-analysis allows us to investigate how linear measurements of gene/protein activity project to the quadratic space of protein interactions in order to re-wire the underlying interactome in each human tissue.

Figure 4 presents the final network. In this network, each node represents a human tissue/cell type and each weighted edge illustrates the extent of overall transcriptional similarity between pairs of tissues. This network is filtered to include only tissue pairs with the highest overlap with each other. In order to assign color to each node, we use *z*-score normalization on the log-transformed alignment mixed *p*-values. Green and red nodes correspond to the highly positive and highly negative range of *z*-scores, which represent similar and dissimilar tissues to yeast, respectively.

**Fig. 4.**
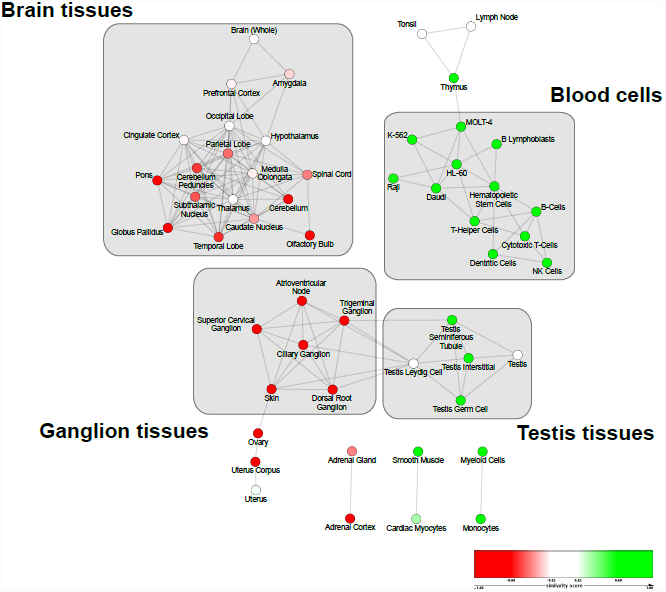
Projection of alignment *p*-values on the network of tissue-tissue similarities. Each node represents a human tissue and edges represent the overall transcriptional similarity among them. Color intensity of nodes represents the similarity/dissimilarity of each tissue to yeast interactome, with colors green and red corresponding to similar and dissimilar tissues, respectively. Group of similar tissues with coherent *p*-values are marked and annotated in the network, accordingly.

Preliminary analysis of this network indicates that the alignment *p*-value of tissues highly correlates with their overall transcriptional overlap. Furthermore, these high-level interactions coincide with each other and fall within distinct *groups* with consistent patterns. We manually identified four such groups and separately annotated them in the network. These groups correspond to brain tissue, blood cells, ganglion tissues, and testis tissues. Among these groups, blood cells and testis tissues exhibit consistent similarity with yeast, whereas brain and ganglion tissues bear consistent dissimilarity.

The existence of homogenous group of tissues with consistent similarity with yeast suggests an underlying conserved machinery in these clusters. This raises the question of what is consistently aligned within each tissue group and how it relates to the computed alignment *p*-values? We address this question, and relate it to the onset of tissue-specific pathologies in the remaining subsections.

### Dissecting tissue-selective genes with respect to their conservation

In this subsection, we investigate the subset of non-housekeeping genes in each homogenous group of human tissues and partition them into sets of genes, and their corresponding pathways that are either conserved in yeast or are human-specific. Next, we analyze how these pathways contribute to the overall similarity/ dissimilarity of human tissues with yeast.

Figure 5 presents the probability density function for the membership distribution of non-housekeeping genes in different human tissues. The observed bi-modal distribution suggests that most non-housekeeping genes are either expressed in a very few selected tissues or in the majority of human tissues. We use this to partition the set of expressed non-housekeeping genes, with the goal of identifying genes that are selectively expressed in each group of human tissues.

**Fig. 5.**
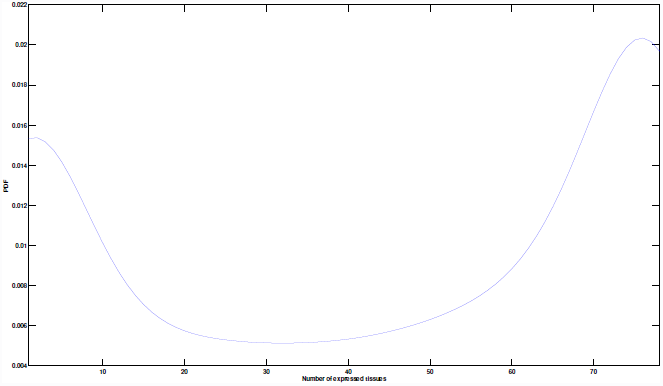
Membership distribution of non-housekeeping genes in human tissues. Number of tissues in which non-housekeeping genes are expressed in is smoothed using normal kernel density to estimated the pdf function. The observed bi-modal distribution suggests that most non-housekeeping genes are either expressed in a very few selected tissues or in the majority of human tissues.

We start with all *expressed non-housekeeping genes* in each tissue group, i.e., genes that are expressed in *at least* one of the tissue members. Next, in order to identify the subset of expressed genes that are *selectively* expressed in each group, we use the *tissue-selectivity* p-*value* of each gene. In this formulation, a gene is identified as selectively expressed if it is expressed in a significantly higher number of tissues in the given group than randomly selected tissue subsets of the same size (see Materials and Methods section for details). Figure 6 illustrates the distribution of tissue-selectivity *p*-values of expressed genes with respect to four major groups in Figure 4. Each of these plots exhibit a bi-modal characteristic similar to the membership distribution function in Figure 5. This can be explained by the fact that membership distribution is a mixture distribution, with the underlying components being the same distribution for the subset of genes that are expressed in different tissue groups. We use critical points of the *p*-value distributions to threshold for tissue-selective genes. The motivation behind our choice is that these points provide shifts in the underlying distribution, from tissue-selective to ubiquitous genes. Given the bi-modal characteristics of these distributions, they all have three critical points, the first of which we use as our cutoff point. This provides highest precision for declared tissue-selective genes, but lower recall than the other two choices.

**Fig. 6.**
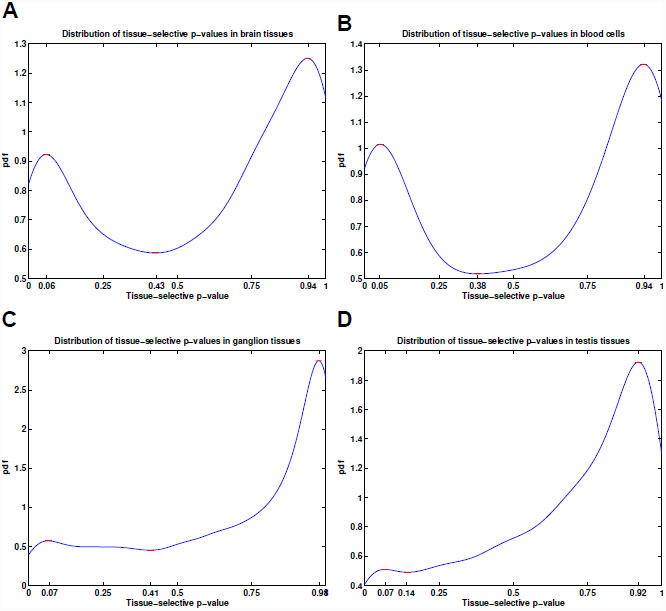
Distribution of tissue-selectivity *p*-values in different tissue groups. (**A**) Brain tissues, (**B**) Blood cells, (**C**) Ganglion tissues, (**D**) Testis tissues. Each plot resembles the same bi-modal distribution as the gene-tissue membership density, with blood cells and brain tissues presenting the most clear separation of tissue-selective genes. The critical points of each distribution function, where the derivative of pdf function is approximately zero, is marked on each plot. These points provide optimal cutoff points for the tissue-selectivity *p*-values as they mark the points of shift in the underlying distribution function.

Having identified the subset of tissue-selective genes with respect to each tissue group, we use the majority voting scheme to tri-partition these sets based on their alignment consistency with yeast. Similar to the procedure we used to tri-partition housekeeping genes, we tried different choices of consensus rate parameter from 50% to 100% with increments of 5%. The percent of unclassified genes decreases linearly with the consensus rate, while relative portions of human-specific/ conserved genes remain the same. We chose 90% for our final results, as it leaves the least number of genes as unclassified, while keeping human-specific and conserved genes well-separated. The set of all tissue-specific genes is available for download as Additional file 1.

Table 3 presents the number of expressed genes, selectively expressed genes, and the percent of tissue-selective genes that are conserved, human-specific, or unclassified within each group of tissues. There is a similar relationship between the ratio of conserved/human-specific genes within each group of tissues and their alignment *p*-values, suggesting that alignment *p*-values are highly correlated with the conservation of tissue-selective genes and their corresponding pathways. Figure 7 illustrates the relative sizes of each subset of genes identified in this study.

**Table 3.** Summary of tissue-selective gene partitioning. CG: Conserved gene, HS: Human-specific gene

| Cluster name | # expressed genes | # TS genes | # CG (%) | # HS (%) | # unclassified (%) |
| --- | --- | --- | --- | --- | --- |
| Brain Tissues | 5936 | 891 | 273 (30.64 %) | 401 (45.01 %) | 217 (24.35 %) |
| Blood Cells | 6092 | 1093 | 460 (42.09 %) | 385 (35.22 %) | 248 (22.69 %) |
| Testis Tissues | 5358 | 328 | 119 (36.28 %) | 126 (38.41 %) | 83 (25.30 %) |
| Ganglion Tissues | 5278 | 274 | 76 (27.74 %) | 136 (49.64 %) | 62 (22.63 %) |

**Fig. 7.**
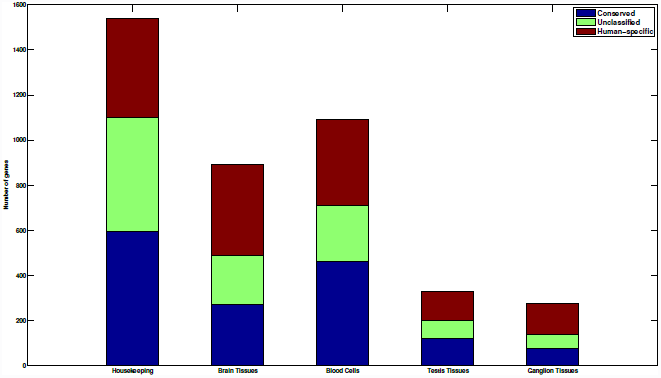
Summary of gene classifications in this study. Housekeeping and tissue-selective genes, in four main groups of human tissues, are classified into three main classes based on their conservation in yeast.

Conserved genes and their corresponding pathways comprise the functional subspace in which we can use yeast as a suitable model organism to study tissue-specific physiology and pathophysiology. On the other hand, human-specific genes provide a complementary set that can be used to construct *tissue-engineered* humanized yeast models. They also provide promising candidates for tissue-specific gene therapies in a similar fashion to NDI1 therapy, in cases where an alternative functional mechanism can be found in yeast. To further investigate these subsets, we focus on blood cells and brain tissues, which illustrate the clearest separation between their tissue-selective and conserved genes in their TSS distribution, and subject them to more in depth functional analysis in next subsections.

### Elucidating functional roles of the brain and blood selective genes

We use g:ProfileR on both human-specific and conserved genes to identify their enriched functions. The complete list of enriched functions is available for download as Additional file 1. These two subsets share many common terms, due to the underlying prior of both being subsets of tissue-selective genes. To comparatively analyze these functions and rank them based on their human-specificity, we use the log of *p*-value ratios between human-specific and conserved genes to filter terms that are at least within 2-fold enrichment. We focus on GO biological processes, KEGG pathways, and CORUM protein complexes and remove all genesets with more than 500 genes to filter for overly generic terms. Finally, to group these terms together and provide a visual representation of the functional space of genes, we use EnrichmentMap (EM)(69), a recent Cytoscape(70) plugin, to construct a network (map) of the enriched terms. We use the log ratio of *p*-values to color each node in the graph. Figures 8 and 9 illustrate the final enrichment map of unique human-specific and conserved blood-selective and brain-selective functions, respectively.

**Fig. 8.**
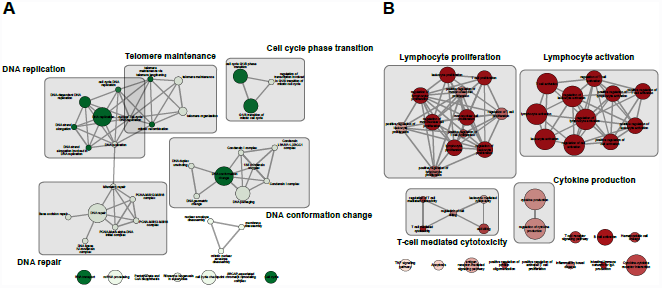
Enrichment map of unique blood-selective functions.

**Fig. 9.**
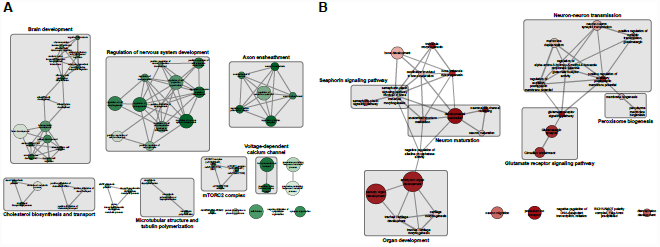
Enrichment map of unique brain-selective functions.

Conserved blood-selective functions, shown in sub-figure ??, are primarily enriched with terms related to DNA replication, cellular growth, and preparing cell for cell-cycle. Among these terms, DNA replication-is tightly linked to both DNA repair and telomere maintenance related terms. Telomere maintenance, specially via telomerase enzyme, is one of the cellular functions that is known to be conserved in yeast, but only active in a selected subset of differentiated human tissues and cell types, including hematopoietic stem cells and male reproductive tissues (71). Functional terms involved in DNA conformation changes, including condensin complex, as well as cell cycle phase transition, specifically from G1 to S phases, are two other groups of conserved functional terms that are highly conserved from yeast to human. On the other hand, human-specific blood-selective functions, shown in Figure ??, are mainly involved in lymphocyte proliferation and activation. Terms in these two groups are also tightly related to each other and form a larger cluster together. In addition, cytokine production and T-cell mediated cytotoxicity also exhibit human-specific, blood-selective characteristics. This is partially expected as these functions are highly specialized immune-cell functions that are evolved particularly in humans to ensure his survival in less-favorable conditions.

Figure ?? shows the functional space of conserved brain-selective functions. Many of these terms correspond to various aspects of brain development, including olfactory bulb, telencephalon, pallium, and cerebral cortex development, as well as the regulatory circuit that controls nervous system development. Considering the unicellular nature of yeast, the exact mechanisms in which orthologs of these pathways modulate yeast cellular machinery is less studied. An in-depth analysis to identify matching phenologs can help us use yeast to study various disorders related to brain development. Another functional aspect that exhibits high conservation is the mTOR complex 2. The target of rapamycin (TOR) signaling is a highly conserved pathway, which forms two structurally distinct protein complexes, mTORC1 and mTORC2. The former complex has a central role in nutrient-sensing and cell growth, and as such, has been used extensively to study calorie restriction (CR) mediated lifespan extension. On the other hand, mTORC2 has been recently proposed to modulate consolidation of long-term memory(72). Cholesterol biosynthesis and transport is another conserved functional aspect that differs significantly from other human tissues. As the most cholesterol-rich organ in the body, expression of genes corresponding to lipoprotein receptors and apolipoproteins is tightly regulated among different brain cells and plays an important role in normal brain development. Dysregulation of these metabolic pathways is implicated in various neurological disorders, such as Alzheimer’s disease(73).Finally, microtubular structure and tubulin polymerization also shows significant conservation and is known to play a key role in brain development(74). These cytoskeletal proteins have recently been associated with brain-specific pathologies, including epilepsy(75).

Finally, we study human-specific brain functions, which are shown in Figure ??. One of the key functional aspects in this group is the semaphorin-plexin signaling pathway. This pathway was initially characterized based on its role in the anatomical structure maturation of the brain, specifically via the repulsive axon guidance, but later was found to be essential for morphogenesis of a wide range of organ systems, including sensory organs and bone development(76). Another human-specific signaling pathway identified in brain is the glutamate receptor signaling pathway, which also cross-talks with circadian entrainment, as well as neuron-neuron transmission. This pathway plays a critical role in neural plasticity, neural development and neurodegeneration(77). It has also been associated with both chronic brain diseases, such as schizophrenia, as well as neurodegenerative disorders, such as Alzheimer’s disease(78).

Both conserved and human-specific genes play important roles in tissue-specific pathologies. In addition, these genes, which are enriched with regulatory and signaling functions, cross-talk with housekeeping genes to control cellular response to various factors. As such, a complete picture of disease onset, development, and progression can only be achieved from a systems point of view. From this perspective, we study not only the genes (or their states) that are frequently altered in disease, but also the underlying tissue-specific and housekeeping pathways in which they interact to exhibit the observed phenotype(s). In the next subsection, we further investigate this hypothesis. We study the potential of different subsets of the identified tissue-selective genes for predicting tissue-specific pathologies.

### Assessing the significance of tissue-specific pathologies among conserved and human-specific tissue-selective genes

To further study the predictive power of tissue-selective genes for human pathologies, we use the *genetic association database* (*GAD*) disease annotations as our gold standard (79). This database collects gene-disease associations from genetic association studies. Additionally, each disease has been assigned to one of the 19 different disease classes in GAD database. We use DAVID functional annotation tool for disease enrichment analysis of tissue-selective genes (80).

First, we seek to identify which disease classes are significantly enriched among each set of tissue-selective genes. Table 4 shows the disease classes enriched in each group of brain and blood selective genes. Conserved blood-selective genes are predominantly enriched with cancers, whereas human-specific blood-selective genes are mainly associated with immune disorders. This can be linked to our previous results indicating that conserved subset is mainly involved in regulating growth, DNA replication, and cell cycle, whereas human-specific genes are primarily involved in lymphocyte proliferation and activation. Conversely, brain-selective genes show higher similarities in terms of disease classes that they can predict. Both conserved and human-specific brain-selective genes can predict psychiatric disorders, but human-specific subset seems to be a more accurate predictor. On the other hand, neurological disorders are only enriched in human-specific subset of brain-selective genes, whereas disorders classified as pharmacogenomic and chemdependency show higher enrichment in conserved genes.

**Table 4.** Enriched disease classes of tissue-selective genes.

|  | Conserved genes |  | Human-specific genes |  |
| --- | --- | --- | --- | --- |
|  | Disease class | *p*-value | Disease class | *p*-value |
| Blood cells | Cancer | $9.29 * 10^{-4}$ | Immune | $1.19 * 10^{-5}$ |
| Brain tissues | Psych | $3.59 * 10^{-4}$ | Psych | $5.70 * 10^{-8}$ |
| | Chemdependency | $2.60 * 10^{-3}$ | Neurological | $2.97 * 10^{-2}$ |
| | Pharmacogenomic | $9.74 * 10^{-2}$ | | |

To summarize the specific disorders that are enriched in each subset of brain-selective genes, we integrate all identified diseases and rank them based on their enrichment *p*-value, if it is only enriched in one set, or their most significant *p*-value, if it is enriched in both sets. Table 5 shows the top 10 disease terms enriched in either human-specific or conserved brain-selective genes. In majority of cases, human-specific genes are more significantly associated with brain-specific pathologies than conserved genes. In addition, there are unique disorders, such as schizophrenia, bi-polar disorder, and seizures, that are only enriched among human-specific genes.

**Table 5.** Comparative analysis of brain-specific pathologies. Top 10 Enriched disorders were identified based on the GAD annotations for conserved and human-specific genes in the brain.

| Disorder | Conserved genes | Human-specific genes |
| --- | --- | --- |
| schizophrenia | 0.008573 | 8.4905E-06 |
| autism | 0.048288 | 0.00077448 |
| dementia | 0.0014356 | - |
| schizophrenia; schizoaffective disorder; bipolar disorder | - | 0.0021433 |
| myocardial infarct; cholesterol, HDL; triglycerides; atherosclerosis, coronary; macular degeneration; colorectal cancer | 0.0051617 | - |
| epilepsy | 0.071562 | 0.0064716 |
| seizures | - | 0.020381 |
| bipolar disorder | 0.048288 | 0.022016 |
| attention deficit disorder conduct disorder oppositional defiant disorder | 0.032444 | 0.023865 |

In conclusion, both conserved and human-specific subsets of tissue-selective genes are significantly associated with different human disorders. However, the human-specific subset shows higher association with tissue-specific pathologies. To this end, they can provide hypotheses for the most appropriate molecular constructs (gene insertions) in yeast to explore molecular/functional mechanisms that cause tissue-specific dysfunction. Such mechanisms can be tested in humans and if validated then yeast can serve as an experimental model for further investigations of biomarkers and pharmacological and genetic interventions.

## Conclusions

In this study, we demonstrated a novel methodology for aligning tissue-specific interaction networks with the yeast interactome and assess their statistical significance. We demonstrated that these alignments can be used to dissect tissue-specific networks into their core component and tissue-specific components. Tissue specific components were used for multiple purposes: (i) by showing that a number of pathologies manifest themselves in dysregulated genes in the tissue-specific group, we motivate exploration of these genes as particularly suitable candidates as drug targets; (ii) by quantifying the alignment of tissue-specific components with yeast, we quantify the suitability of yeast as a model organism for studying corresponding disease/ phenotype; (iii) in cases where there is (statistically) insignificant alignment, it is still possible to use yeast as a model organism, if the dysregulated pathways are aligned; and (iv) in cases where none of these conditions hold, our alignments provide mechanisms for assessing the feasibility of different molecular constructs (gene insertions) for creating more appropriate, tissue-specific, humanized yeast models.

## Materials and methods

### Datasets

#### Protein-protein interaction (PPI) networks

We adopted human tissue-specific networks from *Bossi et al*. (57). They integrated protein-protein interactions from 21 different databases to create the whole human interactome consisting of 80,922 interactions among 10,229 proteins. Then, they extracted the set of expressed genes in each tissue from GNF Gene Atlas and used it to construct the tissue-specific networks, defined as the vertex-induced subgraphs of the entire interactome with respect to the nodes corresponding to the expressed genes in each tissue.

Additionally, we obtained the yeast interactome from the BioGRID(81) database, update 2011(82), version 3.1.94, by extracting all physical interactions, excluding interspecies and self interactions. This resulted in a total of 130,483 (76,282 non-redundant) physical interactions among 5,799 functional elements in yeast (both RNA and protein). Next, we downloaded the list of annotated CDS entries from the Saccharomyces Genome Database (SGD)(83) and restricted interactions to the set of pairs where both endpoints represent a protein-coding sequence, i.e., protein-protein interactions. The final network consists of 71,905 interactions between 5,326 proteins in yeast and is available for download as Additional file 1.

#### Protein sequence similarities between yeast and humans

We downloaded the protein sequences for yeast and humans in FASTA format from Ensembl database, release 69, on Oct 2012. These datasets are based on the GRCh37 and EF4 reference genomes, each of which contain 101,075 and 6,692 protein sequences for *H. Sapiens* and *S. Cerevisiae*, respectively. Each human gene in this dataset has, on average, 4.49 gene products (proteins). We identified and masked low-complexity regions in protein sequences using *pseg* program(84). The *ssearch36* tool, from *FASTA*(85) version 36, was then used to compute the local sequence alignment of the protein pairs using the Smith-Waterman algorithm(86). We used this tool with the BL0SUM50 scoring matrix to compute sequence similarity of protein pairs in humans and yeast. All sequences with E-values less than or equal to 10 are recorded as possible matches, which results in a total of 664,769 hits between yeast and human proteins. For genes with multiple protein isoforms, coming from alternatively spliced variants of the same gene, we only record the most significant hit. The final dataset contains 162,981 pairs of similar protein-coding genes, and is available for download as Additional file 1.

### Sparse network alignment using belief propagation

Analogous to the sequence alignment problem, which aims to discover conserved genomic regions across different species, network alignment is motivated by the need for extracting shared functional pathways that govern cellular machinery in different organisms. The network alignment problem in its abstract form can be formulated as an optimization problem with the goal of identifying an optimal mapping between the nodes of the input networks, which maximizes both sequence similarity of aligned proteins and conservation of their underlying interactions. At the core of every alignment method are two key components: i) a scoring function and ii) an efficient search strategy to find the optimal alignment. The scoring function is usually designed to favor the alignment of similar nodes, while simultaneously accounting for the number of conserved interactions between the pair of aligned nodes. Biologically speaking, this translates to identifying functional *orthologs* and *interologs*, respectively.

Given a pair of biological networks, *G* = (*V*_*G*_, *E*_*G*_) and *H* = (*V*_*H*_, *E*_*H*_), with *n*_*G*_ = |*V*_*G*_| and *n*_*H*_ = |*V*_*H*_| vertices, respectively, we can represent the similarity of vertex pairs between these two networks using a weighted bipartite graph *L* = (*V*_*G*_ * *V*_*H*_, *E*_*L*_, *w*), where *w* : *E*_*L*_ → *ℛ* is a weight function defined over edges of *L*. We will denote mapping between vertices *v*_*i*_ ∈ *V*_*G*_ and *v*_*i'*_ ∈ *V*_*H*_ with (*i, i′*) and *ii′*, interchangeably. Let us encode the edge conservations using matrix 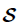, where 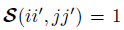, iff alignment of *v_i_* → *v_i′_* together with *v_j_* → *v_j′_* will conserve an edge between graphs *G* and *H*, and 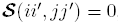, otherwise. Then, the network alignment problem can be formally represented using the following integer quadratic program:

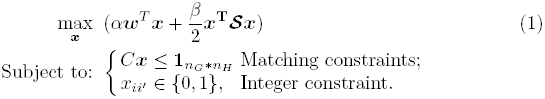

Here, *C* is the incidence matrix of graph *L* and *x* is the matching indicator vector. In this formulation, the *weight* of each alignment, 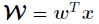, accounts for the similarity of aligned nodes, while the alignment *overlap*, 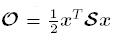, counts the number of conserved edges under the given alignment. When *L* is a complete bipartite graph, each pair of vertices between *G* and *H* represents a viable candidate, in which case we have 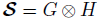. However, *Bayati et al*. (59) recently proposed an efficient method, based on the message passing algorithm, for cases where *L* is sparse, i.e., |*E_L_*| << *n*_*G*_ * *n*_*H*_, by restricting the search space to the subset of promising candidates that are provided by *E*_*L*_. We use this algorithm for solving the network alignment problem.

### Tissue-specific random model (TRAM) for generating pseudo-random tissues

Let us denote the global human interactome by *G* = (*V*_*G*_, *E*_*G*_) and each tissue-specific network by *G*_*T*_ = (*V*_*T*_, *E*_*T*_), respectively. using this notation, we have *n*_*T*_ = |*V*_*T*_|, *V*_*T*_ ⊂ *V*_*G*_, and *E*_*T*_ ⊂ *E*_*G*_ is the subset of all edges from *G* that connect vertices in *V*_*T*_, i.e., *G*_*T*_ is the vertex-induced subgraph of *G* under *V*_*T*_. We note that the existence of universally expressed genes, corresponding to housekeeping proteins, enforces a unique topology for human tissue-specific networks with a shared, dense core connected to peripheral tissue-specific proteins. We propose a new random model for explicitly taking advantage of this prior knowledge and create pseudo-random networks that respect the underlying topology of tissue-specific networks. Let *V*_*U*_ denote the set of universally expressed genes across all human tissues, having size *n*_*U*_ = |*V*_*U*_|. Moreover, let *G*_*U*_ = (*V*_*U*_, *E*_*U*_) be the shared, vertex-induced subgraph imposed by housekeeping genes. We create an ensemble of *pseudo-random tissues* for each given *G*_*T*_, denoted by 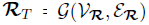, in two steps. First, we sample pseudo-random vertex sets of size *n*_*T*_, denoted by 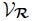, by fixing all the housekeeping genes, *V*_*U*_, and sampling *n*_*T*_ − *n*_*U*_ vertices without replacement from *V*_*G*_ − *V*_*U*_. Then, we construct the pseudorandom ensemble 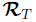 as the set of vertex induced subgraphs of *G* imposed by 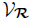.

### Significance of network alignments

For each optimal alignment of a human tissue-specific network with yeast, given by its indicator variable *x*, we quantify the overall sequence similarity of aligned proteins with the matching score of the alignment, 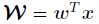, and the total number of conserved edges by the alignment overlap, 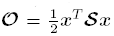. These measures can be used to rank different network alignments. However, without a proper reference to compare with, it is almost impossible to interpret these values in a statistical sense. To address this, we use the *pseudo-random tissue* model as our random model and empirically compute a *topological*, a *homological* (sequence-based), and a *mixed* alignment *p*-value for each alignment using a Monte-Carlo simulation. To this end, we sample *k*_𝓡_ pseudo-random tissues of the same size and align each of them separately with yeast.

Let 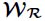 and 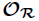 be the random vectors representing the weight and overlap of aligning random tissues with yeast, respectively. First, we define individual *p*-values for the conservation of network topology and sequence homology. Let us denote by 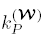 and 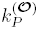 the number of random samples that have weight and overlap greater than or equal to the original alignment, respectively. Then, we can define the following *p*-values:

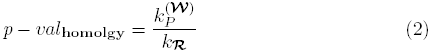

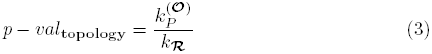

Before we define the *mixed* p-value, we define an upper bound and a lower bound on the p-value that is independent of the mixing function. For cases where both 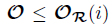 and 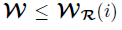, for 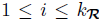, we can report that the random alignment is at least as good as the original alignment. Conversely, if both 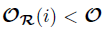 and 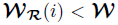, we can assert that the original alignment outperforms the random alignment. Let us denote the number of such cases by *k*_*P*_ and *k*_*N*_, respectively. Using this formulation, we can compute the following bounds on the mixed p-value of the alignment:

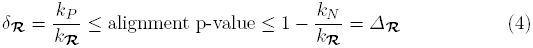

We can use these bounds to estimate the similarity of each tissue-specific network to the yeast interactome. Tissues where the upper-bound of the alignment p-value is smaller than a given threshold *α*_*u*_ are considered similar to yeast, while tissues with the lower-bound larger than *α*_*l*_ are considered dissimilar. We note that the cases with contradictory results for weight and overlap, either if 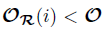 and 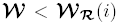, or 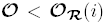 and 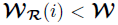, are not straightforward to interpret. To quantify this ambiguity, we define the confidence of a p-value interval as 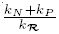. Finally, we define a mixed p-value based on the mixing function of the network alignment. Let us define a new random variable 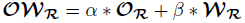. Finally, we define the mixed p-value as:

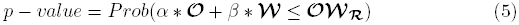

### Differential expression of genes with respect to a group of tissues

Given a homogenous group of human tissues/cell types, we first identify all *expressed genes* in the group, i.e., all non-housekeeping genes that are expressed in *at least* one of the tissue members. Next, in order to identify the subset of expressed genes that are *selectively* expressed, we use a *hypergeometric* random model. A gene is identified as selectively expressed if it is expressed in significantly higher number of tissues in the given group than randomly selected tissue subsets of the same size. Let *N* and *n* denote the total number of tissues in this study and the subset of tissues in the given group, respectively. Moreover, let us represent by *c*_*N*_ the number of all tissues in which a given gene is expressed, whereas *c*_*n*_ similarly represents the number of tissues in the given group that the gene is expressed. Finally, let the random variable *X* be the number of tissues in which the gene is expressed, if we randomly select subsets of tissues of similar size. Using this formulation, we can define the *tissue-selectivity* p-*value* of each expressed gene in the given group as follows:

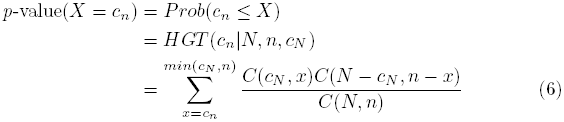

In order to partition genes into *selective* and *ubiquitous* genesets, we derive the tissue-selectivity *p*-value distribution of all expressed non-housekeeping genes in the given group. We use the Gaussian kernel to smooth this distribution and then find the critical points of the smoothed density function to threshold for tissue-selective genes. The motivation behind our choice is that these points provide shifts in the underlying distribution, from tissue-selective to ubiquitous genes. Given the bi-modal characteristic of the distribution, it has three expected critical points. We use the first of these points as our cutoff point. This provides highest precision for declared tissue-selective genes, but lower recall than the other two choices.

### Conservation of genesets based on the majority voting rule

Given a set of genes that are selectively expressed in a homogenous group of tissues/cell types, we are interested in tri-partitioning them into either *conserved*, *human-specific*, or *unclassified* genes. *Conserved genes* are the subset of tissue-selective genes that are consistently aligned in majority of aligned tissues in the given group. Conversely, *human-specific genes* are the subset of tissue-selective genes that are consistently unaligned in majority of tissues in the given group. Finally, *unclassified genes* are the subset of tissue-selective genes for which we do not have enough evidence to classify them as either conserved or human-specific.

The key data-structure we use to tri-partition genesets is the *alignment consistency table*. Let *C* be a group of homogenous tissues with *n* = |*C*|. Furthermore, let 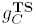 represent the set of tissue-selective genes with respect to *C*, such that 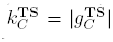. The alignment consistency table is a table of size 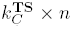, represented by 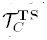, in which 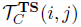 is the aligned yeast partner of *i*^*th*^ tissue selective gene under the network alignment of *j*^*th*^ tissue in *C*, or *′–′* (gap), if it is unaligned. We find the most common partner for each tissue-selective gene and use a *consensus rate*, represented by *τ*, to summarize each rows of the alignment consistency table. If a gene is consistently aligned to the same yeast partner in at least *τ* * *n* tissues in *C*, we declare it as conserved. Similarly, if it is unaligned in at least *τ* * *n* tissues in *C*, we classify it as human-specific. If neither one of these conditions hold, we report it as unclassified.

## Availability of supporting data

The data sets supporting the results of this article are included within the article and its additional files.

## Author’s contributions

SM proposed the initial idea of research, conceived of the study, and prepared the manuscript. SM and BS designed and implemented most of methods and performed the experiments. SS helped with the experimental design, as well as analyzing and interpreting the biological implications of the results. AG provided guidance relative to the theoretical and practical aspects of the methods, and design of proper statistical model(s) to validate the results. All authors participated in designing the structure and organization of final manuscript. All authors read and approved the final manuscript.

## Acknowledgements

This work is supported by the Center for Science of Information (CSoI), an NSF Science and Technology Center, under grant agreement CCF-0939370, and by NSF grants DBI 0835677 and 0641037.

## Additional Files

**Additional file 1 — network alignments**

Compressed (*.zip) file containing individual tissue-specific alignments.

**Additional file 2 — HK genes**

List of housekeeping genes and their classifications into conserved, human-specific, and unclassified subsets

**Additional file 3 — Core gene alignment**

Alignment graph of core housekeeping genes

**Additional file 4 — HK Enrichment**

Functional enrichment analysis of different subsets of HK genes

**Additional file 5 — Alignment statistics**

Alignment statistics for each tissue alignment

**Additional file 6 — TS genes**

Tissue-selective genesets and their respective classifications for brain tissues, blood cells, testis tissues, and ganglion tissues

**Additional file 7 — TS Enrichment**

Functional analysis of different subsets of tissue-selective genes

**Additional file 8 — PPI Nets**

Protein-protein interaction networks used as input in this study.

**Additional file 9 — Sequence similarities**

Sequence similarity between yeast and human proteins.

